# Participant engagement and feedback in microbiome projects: a case of AWI-Gen 2

**DOI:** 10.64898/2026.04.20.718838

**Authors:** Claudine Nkera-Gutabara, Luicer A. Ingasia Olubayo, Ovokeraye O. Oduaran, Isaac Kisiangani, Simon Khoza, Khulu Gama, Meriam Maritze, Carol Mabunda, Dorcas Keya, Kayode E. Adetunji, Stephen Tollman, Lisa K. Micklesfield, Shukri F Mohamed, F Xavier Gómez-Olivé, Furahini Tluway, Michèle Ramsay, Ami S Bhatt, Scott Hazelhurst, Dylan G Maghini, the AWI-Gen Collaborative Centre, MADIVA Research Hub

## Abstract

Returning individualized microbiome results in ways that are ethical, comprehensible, and useful remains under-explored in African settings. We nested a multi-site, mixed-methods study within the AWI-Gen Wave 2 gut microbiome sub-study of 1,801 women aged 42 – 86 years to engage the participants and provide feedback. All (1,001) participants from Agincourt and Soweto (South Africa) and Nairobi (Kenya) were invited to feedback meetings: 496 from Agincourt, 87 from Soweto, and 195 from Nairobi responded. Engagement strategies were tailored by site (small-group and home-based sessions, visual metaphors, Foldscopes, and local-language delivery). Using semi-structured discussions and structured observations analysed thematically in MAXQDA under COREQ, five cross-cutting themes emerged: (1) understanding of microbiome reports, (2) emotional responses to feedback, (3) perceived health relevance, (4) trust in research institutions, and (5) suggestions for improving engagement. Culturally grounded explanations and local-language facilitation enhanced comprehension and perceived relevance; English-heavy sessions were associated with more confusion. Most participants expressed satisfaction and described planned or enacted dietary and lifestyle changes, while frustration centred on long delays between sampling and feedback. Trust increased with transparency and individualized return of results but was often conditional on minimizing burdensome procedures such as repeat blood sampling (phlebotomy) and ensuring timely feedback. Engagement was feasible and low-cost (approximately USD 29-59 per participant) with site-specific resource needs. Limitations included constrained generalizability beyond the three study sites. Returning individualized microbiome findings in community settings in Africa is acceptable, feasible, and can motivate health-promoting behaviours when delivered promptly and in culturally and linguistically appropriate ways.

**IMPORTANCE:** Microbiome studies rarely return individualized results in low-resource settings due to concerns about appropriate feedback and associated costs. This gap risks eroding trust and diminishing research impact. In three African communities, tailored feedback on gut microbiome profiles was provided to 778 women. By documenting a costed, multi-site engagement model and the themes influencing acceptance and actionability, this work offers a practical framework for ethically returning complex -omics results at scale in underrepresented populations - advancing scientific equity and strengthening community trust in microbiome research.

## INTRODUCTION

Participant engagement in research involving biological samples or personal data is both ethically imperative and operationally essential. It fosters trust, encourages participation in future projects, and ensures that research results and benefits are shared. However, feedback must be meaningful, scientifically accurate, and delivered in ways that minimize potential harm and increase understanding. Data generated from the human microbiome analyses further raises social and ethical questions around ownership, interpretation, and communication of biological data [1, 2].

In large-scale genomics and population-based studies such as those conducted by the AWI-Gen Collaborative Centre [3], engagement typically follows a three-tiered model: (1) community-level consultations prior to recruitment and consent; (2) provision of point-of-care results during data collection (e.g., HIV testing, blood pressure) with counselling or referrals if necessary; and (3) community meetings at study conclusion to share aggregate findings [4]. While returning simple point-of-care results is valued and straightforward, sharing complex findings such as genomic or microbiome profiles is more challenging – scientifically, logistically, and ethically. These challenges have prompted calls to embed social science and ethics perspectives within microbiome research to understand how participants interpret and act on their data [2]. Feedback is therefore not merely a scientific exercise but a relational act shaped by trust, language, and context.

Participatory initiatives such as the American Gut Project and the Isala study [5, 6, 7, 8] illustrate both the promise and the communication hurdles of returning individualized microbiome results. However, these efforts have largely occurred in high-resource, digitally connected settings. In African contexts – where historical exclusion from genomic and microbiome research is being actively addressed – returning results requires heightened sensitivity and an emphasis on reciprocity, transparency, and benefit-sharing [9, 10, 4]. Research from the continent shows that participants strongly prefer to receive personal health information, yet persistent questions remain about what constitutes useful feedback and how it can be ethically delivered [11, 12, 13]. Because microbiome composition is influenced by social determinants such as diet, healthcare access, and socioeconomic status [14], meaningful feedback must address these structural realities. Even if they were appropriate, email or a paper postal system are often not available (and not at our sites).

Returning microbiome results can also promote “scientific citizenship,” positioning participants as collaborators rather than mere data sources. Such engagement can enhance scientific literacy, support informed health decisions, and strengthen trust when feedback is personalized, culturally appropriate, and accessible. The AWI-Gen 2 microbiome sub-study, conducted across urban and rapidly transitioning rural communities in South Africa and Kenya, provides a unique opportunity to examine these processes in under-represented African contexts. Building on prior calls to integrate social-science perspectives into microbiome research [2], we empirically examine how personalized microbiome results are understood and acted upon in everyday life. By returning individual microbiome reports to over 700 women aged 42–86 years and tailoring engagement through visual metaphors (e.g., soil biodiversity, soccer teams), Foldscopes [15], and small-group or home-based sessions, this study explores what participants found meaningful or concerning, and how these interactions shaped perceptions of health, research, and microbiome science.

## MATERIALS AND METHODS

### DESCRIPTION OF STUDY AND SITES

The Africa Wits-INDEPTH Partnership for Genomic Studies (AWI-Gen), part of the H3Africa initiative, is a large-scale genomic and epidemiological initiative investigating genetic and environmental contributions to cardiometabolic diseases in African populations. AWI-Gen spans two waves of data collection: 2013-2017 for Wave 1 and 2018-2022 for Wave 2, across research sites in Burkina Faso, Ghana, Kenya, and South Africa. Data was generated through partnerships with Health and Demographic Surveillance System (HDSS) sites, providing rich demographic, clinical, and biological data[3].

The AWI-Gen 2 microbiome study is a sub-study of the second wave of AWI-Gen which ended in 2022, and randomly sampled 1,801 (out of 7225 in AWI-Gen 2) adult women aged between 42 and 86 years across three sites, Soweto, Agincourt and Nairobi. The microbiome study aimed to characterize gut microbial diversity and its association with health outcomes among middle-aged and older adult women across sub-Saharan Africa [16]. The data collected included stool, blood and urine samples, clinical information and health and lifestyle questionnaire responses. Participants represented diverse lifestyles rarely captured in microbiome studies, including rural farming communities, towns transitioning toward industrialization, and dense urban settlements.

Community engagement and feedback on results were integral to the broader AWI-Gen framework, which included advisory board consultations, return of point-of-care results during data collection, and community-level dissemination of genomic findings. The microbiome sub-study was embedded within this structure, with participants informed at recruitment and later provided microbiome-specific feedback once metagenomic analyses were completed.

In the current pilot study, community engagement and participant results feedback activities were conducted at three of the six sites: MRC/Wits Rural Public Health and Health transitions Research Unit (Agincourt) and Development Pathways for Health Research Unit in Soweto (South Africa), and Nairobi Urban health demographic surveillance systems (Kenya). The formal education levels of the microbiome participants at the sites are shown in Figure **1**.

**FIG 1.**
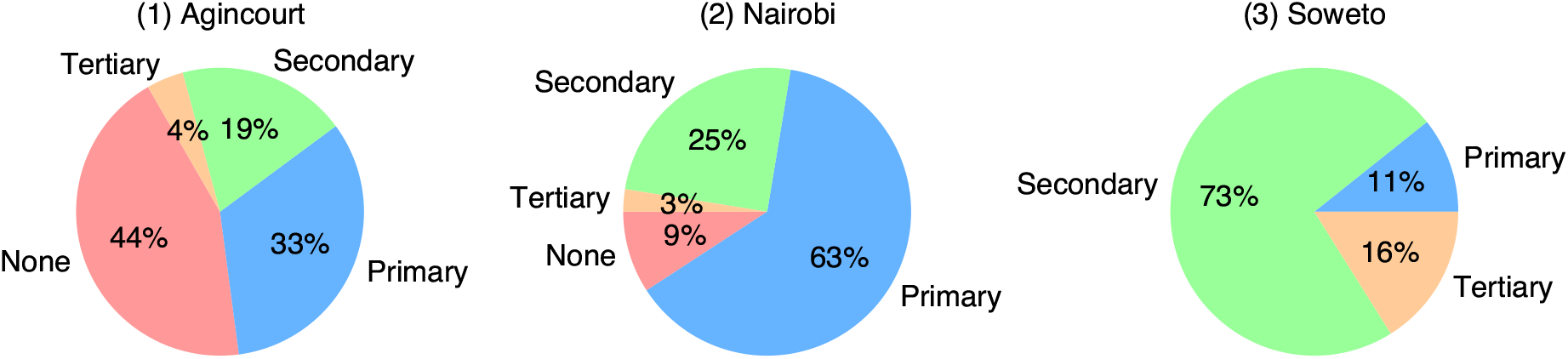
Formal education: Proportion of participants who attained this as their highest level of education (not necessarily completed) at the different sites.

#### Agincourt, South Africa

The HDSS is located in a rural site located in the Bushbuckridge sub-district of Mpumalanga province and is part of the MRC/Wits Rural Public Health and Health Transitions Research Unit. Established in 1992, Agincourt HDSS covers 27 villages and has a long history of migrancy, shaped by forced removals during apartheid and refugee influx from Mozambique in the 1980s. The population is culturally diverse, with high levels of poverty, incomplete infrastructure, and ongoing epidemiological transition (Figure **2**).

**FIG 2.**
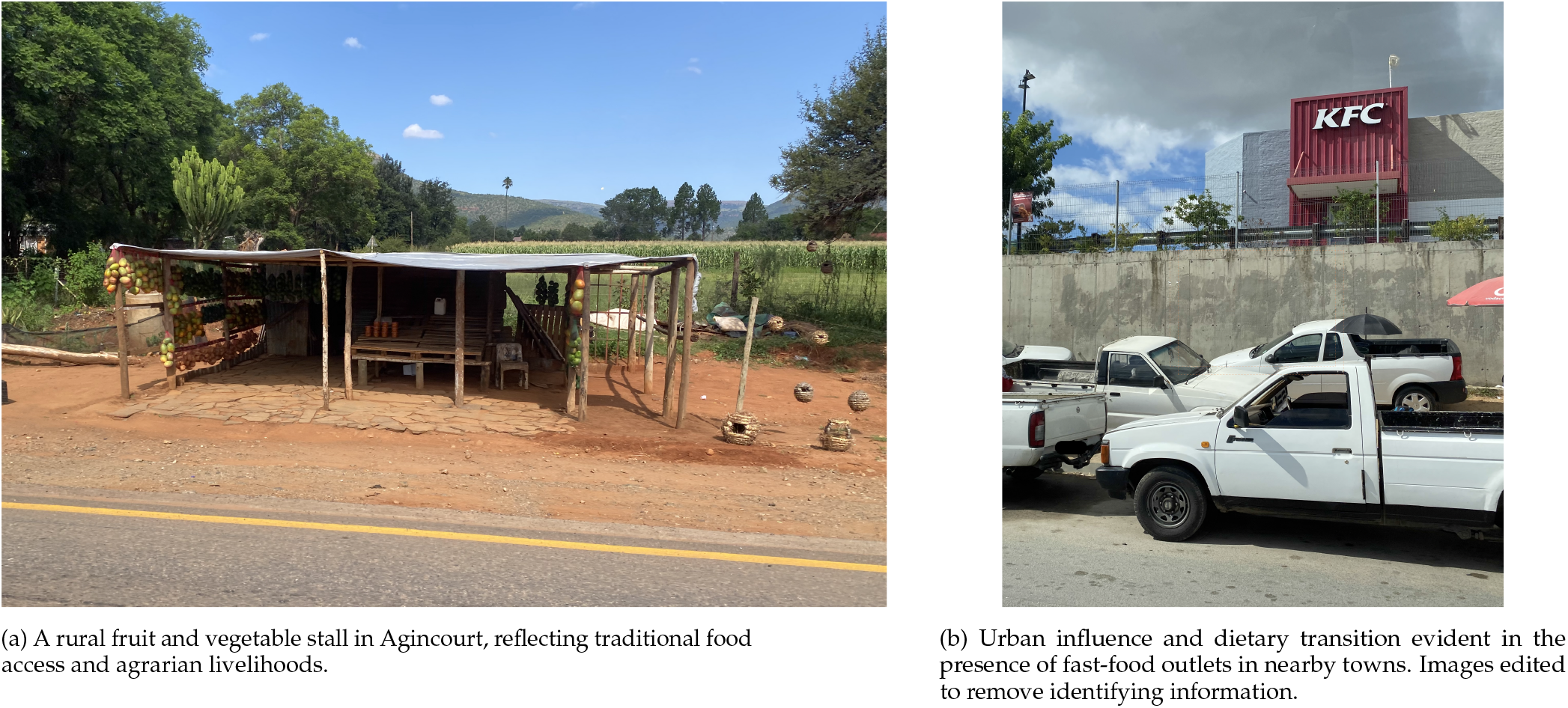
The Agincourt HDSS, Bushbuckridge sub-district of Mpumalanga Province in South Africa.

#### Soweto, South Africa

A historically significant and densely populated urban area in Johannesburg, Soweto is socioeconomically heterogeneous, ranging from informal settlements to middle-class areas. Despite formal inclusion in Johannesburg’s administrative system post-1994, disparities in access to sanitation and public health infrastructure persist, alongside high mobility and internal migration (Figure **3**).

**FIG 3.**
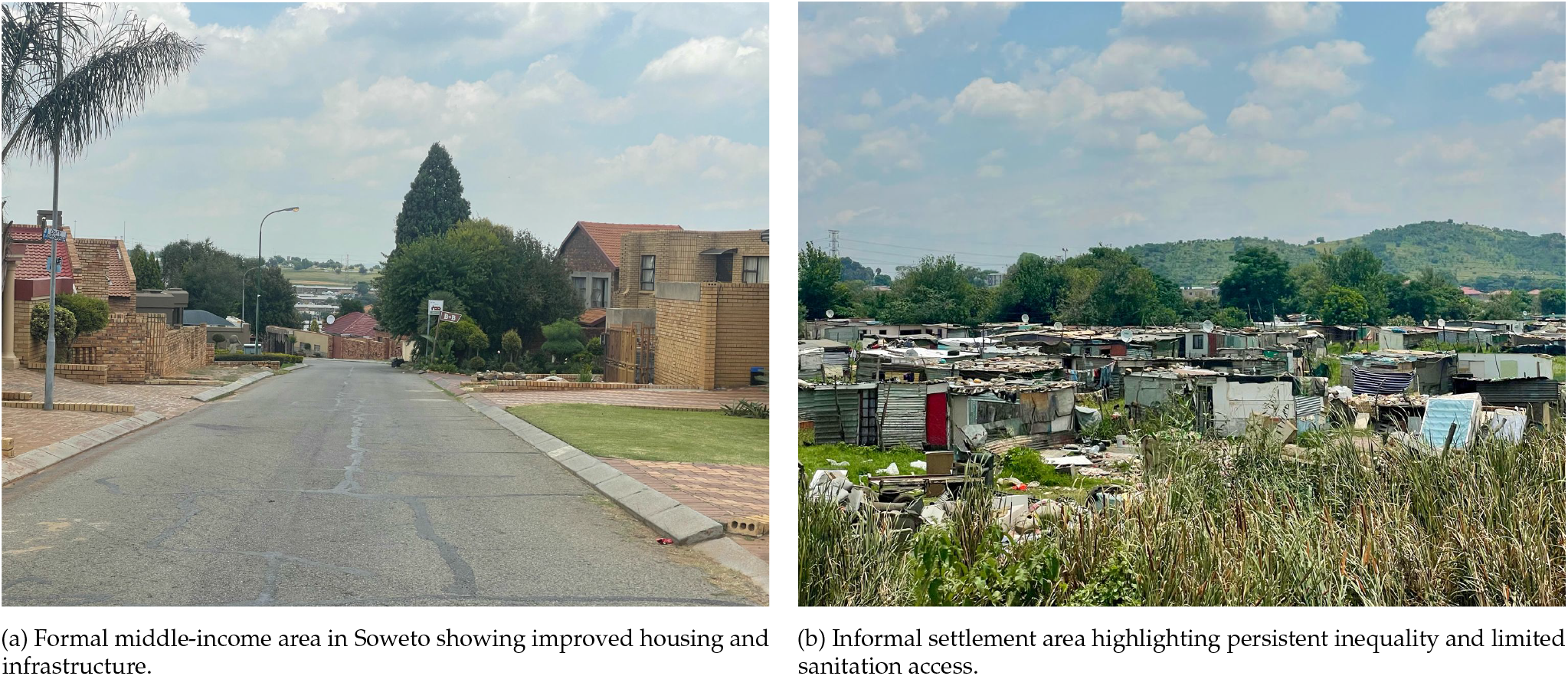
The Soweto Area, Johannesburg, Gauteng Province in South Africa, managed by the Wits Developmental Pathways Research Health Unit within the Chris Hani Baragwanath Academic Hospital.

#### Nairobi, Kenya

The Nairobi Urban HDSS (NUHDSS), established in 2002, covers two informal settlements – Korogocho and Viwandani – home to over 85,000 residents [17]. These communities experience some of the worst socio-economic and health outcomes in Nairobi. Korogocho’s population is more stable, while Viwandani is characterized by higher mobility due to its proximity to industrial zones (Figure **4**).

**FIG 4.**
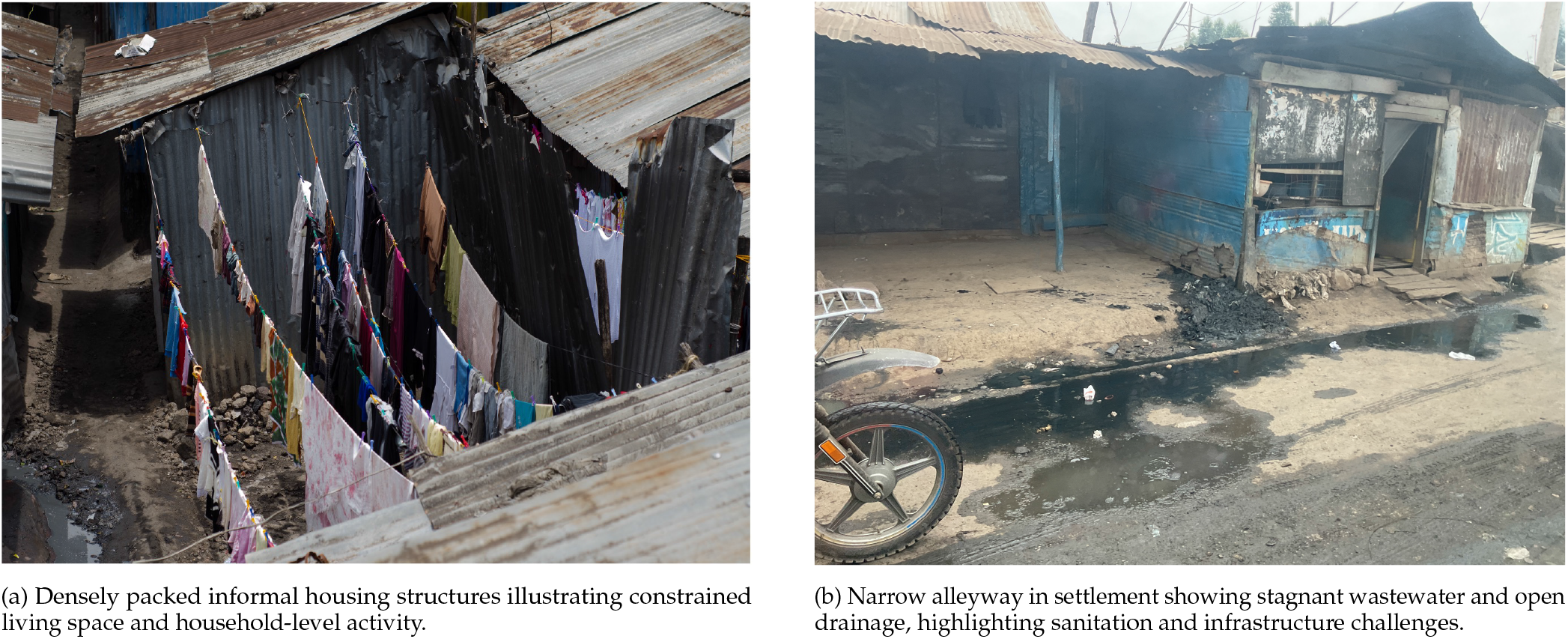
The informal settlements in Nairobi (Viwandani and Korogocho), Kenya, managed by the African Population and Health Research Center (APHRC).

#### Study Design

This was a qualitative, multi-site, community-based study nested within the larger AWI-Gen 2 microbiome sub-study (Figure **5**). The engagement and feedback initiative aimed to assess participant responses to personalised gut microbiome results and to evaluate context-appropriate dissemination strategies. A mixed-methods approach combined systematic documentation of engagement processes with interpretive exploration of participants’ experiences and reflections. This dual approach enabled evaluation of both the effectiveness of engagement and feedback strategies and the depth of participant understanding. Having established the overall design, we next describe the composition and training of the research team, which was central to ensuring reflexive, context-sensitive facilitation. The study followed COREQ (Consolidated Criteria for Reporting Qualitative Research) guidelines [18].

**FIG 5.**
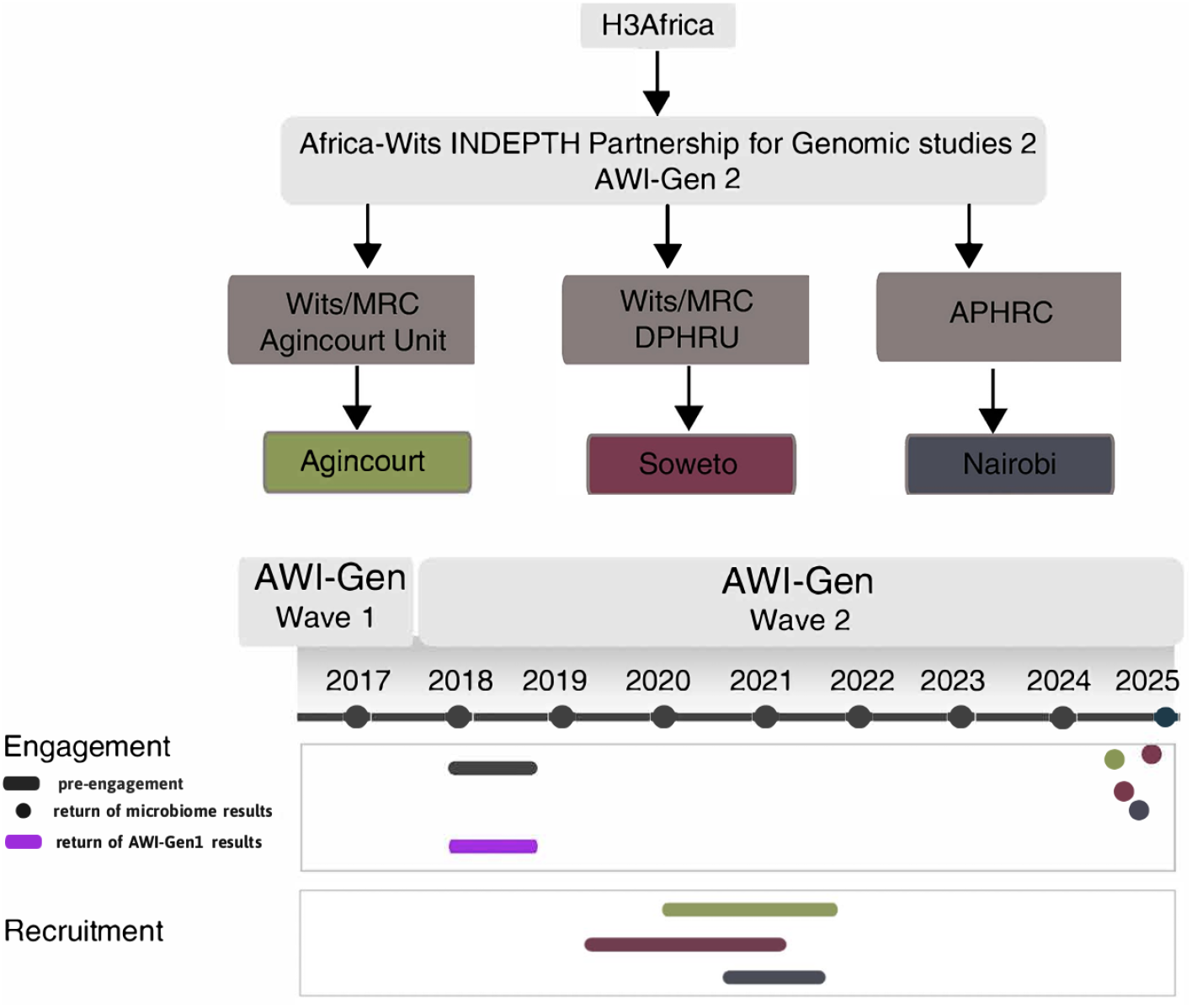
Description of the AWI-Gen study across two waves of data collection. Timeline showing recruitment during Wave 1 and Wave 2 and completed microbiome community engagement.

#### Research team and reflexivity

The research team comprised investigators, community engagement specialist, fieldworkers affiliated with Sydney Brenner Institute for Molecular Bioscience (SBIMB), APHRC (Kenya), MRC/Wits (Agincourt), and MRC/Wits DPHRU (Soweto). Fieldworkers were recruited from the respective communities and trained in qualitative methods, ethics, and reflexive facilitation techniques. Existing relationships with the community and participants via the research site and HDSS platforms promoted cultural congruence and a sense of familiarity, which in turn facilitated open dialogue.

All facilitators used a structured guide and followed standardized templates for observation. Regular team debriefings allowed for reflection on positionality and bias mitigation. Interviews and discussions were conducted in participants’ preferred languages (XiTsonga, IsiZulu, or Kiswahili).

#### Participant selection

The feedback activity invited *all* individuals from the three sites who had previously provided stool samples as part of the AWI-Gen 2 microbiome sub-study (*n*=1,001 across Agincourt (*n*=538), Soweto (*n*=226), and Nairobi (*n*=237)). Our intention was to provide feedback to each microbiome participant available as part of the study’s ethical commitment to return results. The existing updated HDSS and research site records were used to invite participants to receive their individual gut microbiome results. Participants were approached through home visits, phone calls, and community liaisons. After participants had received and discussed their individual microbiome report at the feedback sessions, facilitators proceeded by introducing the purpose of the current study. They obtained verbal and written informed consent to administer open-ended qualitative questions related to understanding and reflections on feedback during the same feedback session.

#### Research instruments

A semi-structured topic guide was collaboratively developed by the core team and community staff, which included four thematic areas: (1) understanding and interpretation of gut microbiome results; (2) perceived value and relevance of the feedback; (3) behavioural reflections on health, anticipated change, and future participation; and (4) suggestions for improving engagement. The guide was translated into local languages and tested for cultural suitability. To support participant comprehension, facilitators used locally familiar metaphors (e.g., gardens, soccer teams) and visual tools such as Foldscopes – low-cost paper microscopes – that enabled participants to view prepared microbial slides and local water samples. A detailed description is provided in Supplemental File S1.

#### Data collection

Following the completion of primary metagenomic analysis and the generation of individual-level gut microbiome profiles, community engagement, participant results feedback, and data collection activities were conducted in May 2024 in Agincourt, July and October 2024 in Soweto and October 2024 in Nairobi. Site teams mobilized participants to disseminate the results through existing HDSS and research site infrastructure, community liaison officers, and previously collected contact records. The mobilization process spanned 2 to 4 weeks per site and involved confirmation of participant availability, tailoring engagement formats to local contexts, and training fieldworkers and facilitators.

Each site adopted a tailored engagement model reflecting its demographic, infrastructure, and sociocultural characteristics. In Agincourt, participants in villages with large numbers attended focused group discussions (FGDs) in community halls, while individual home visits were conducted for participants living in villages with few participants. These personalized discussions usually lasted between 30 - 60 minutes. In Soweto, two large group audio recorded sessions were held in a public venue. Focus was placed on concise, locally translated reports that addressed urban health stressors (e.g., diet, stress, processed food) and how these impact gut health. Due to high mobility, follow-up phone interviews were conducted, transcribed and translated, with select participants (*n*=37) and allowed a deeper one-on-one exploration of reactions and interpretations after the engagement session. In Nairobi, audio recorded engagement was conducted in smaller group sessions (≈10 participants), using interactive, locally grounded metaphors. Feedback sessions were held in community centres. Visual metaphors (e.g., soccer team positions representing microbial diversity and function) were used to enhance understanding in a familiar, non-intimidating way.

Facilitators observed non-verbal communication and recorded field notes using structured templates, with triangulation between transcripts and observations enriching contextual analysis. Data saturation was considered achieved when no new themes emerged across two consecutive FGDs per site. The bias of observation was minimized by training fieldworkers and facilitators to avoid projecting interpretations onto participants behaviours.

#### Data management and analysis

All audio recordings from Nairobi and Soweto were transcribed verbatim and translated into English as needed. In Agincourt, field notes were captured contemporaneously and entered into REDCap. Transcripts and notes were anonymized, stored securely on password-protected servers, and accessed only by the analysis team. Data were imported into MAXQDA 2022 (version 24.9.0) (VERBI Software, 2021) [19] for coding and management.

Thematic analysis combined deductive strategies based on predefined domains of the topic guide with inductive coding to capture new concepts emerging from participants’ narratives. Two analysts independently coded a subset of transcripts to develop an initial frame, after which a shared codebook was iteratively refined and applied to the full dataset. Regular team discussions reconciled differences and finalized the thematic structure, enhancing rigour and reliability. Themes were mapped across sites to identify both shared and context-specific patterns and were organized hierarchically into major themes, minor themes, and subthemes.

Reflexivity was embedded throughout the analysis process. The analysts maintained reflective memos, and debriefings examined how positionality might influence interpretation. Although transcripts were not returned for member checking due to logistical constraints, facilitators summarised key points at the end of each session to enable immediate participant validation.

#### Ethical consideration

All participants provided written informed consent for results feedback and audio-recorded discussions, and additional verbal consent was obtained for follow-up phone interviews in Soweto. Participants were informed of their right to decline participation or withdraw at any time without consequence. Reports were distributed individually or privately or in small-group formats and participants were advised not to share personal health information publicly during group sessions. Particular attention was paid to minimize potential harm or distress when sharing microbiome findings, especially when participants expressed confusion, concern, or feelings of guilt about their gut health profiles. Facilitators were trained to contextualize the findings with sensitivity and emphasize that the results were not diagnostic, but exploratory and educational. Anonymity was ensured during transcription and analysis through de-identification, and transcripts were stored securely. Ethical approval was granted by the University of the Witwatersrand Human Research Ethics Committee and relevant institutional review boards at each study site (Protocol No.: M170880, M2210108).

## RESULTS

A total of **778 participants** across the three sites attended the personalized microbiome feedback sessions (Agincourt, n = 496; Soweto, n = 87; Nairobi, n = 195). The remaining 224 participants did not attend due to reasons such as relocation, work obligations, death and/or outdated contact details.

Analysis of the participant feedback was guided by three open-ended questions:

1. What are the benefits of the results for you?
2. Do these results motivate you to join future studies?
3. Do you have any questions or recommendations?

Thematic analysis yielded five major themes: (1) understanding of microbiome reports; (2) emotional responses to feedback; (3) perceived health relevance; (4) trust in research institutions; and (5) suggestions for improving engagement. These themes, which cut across sites while also reflecting unique contextual nuances, are quantified (Table S2.1) and illustrated with participant quotes provided in Supplemental File S2, and organized with their corresponding subthemes in Figure **6**.

**FIG 6.**
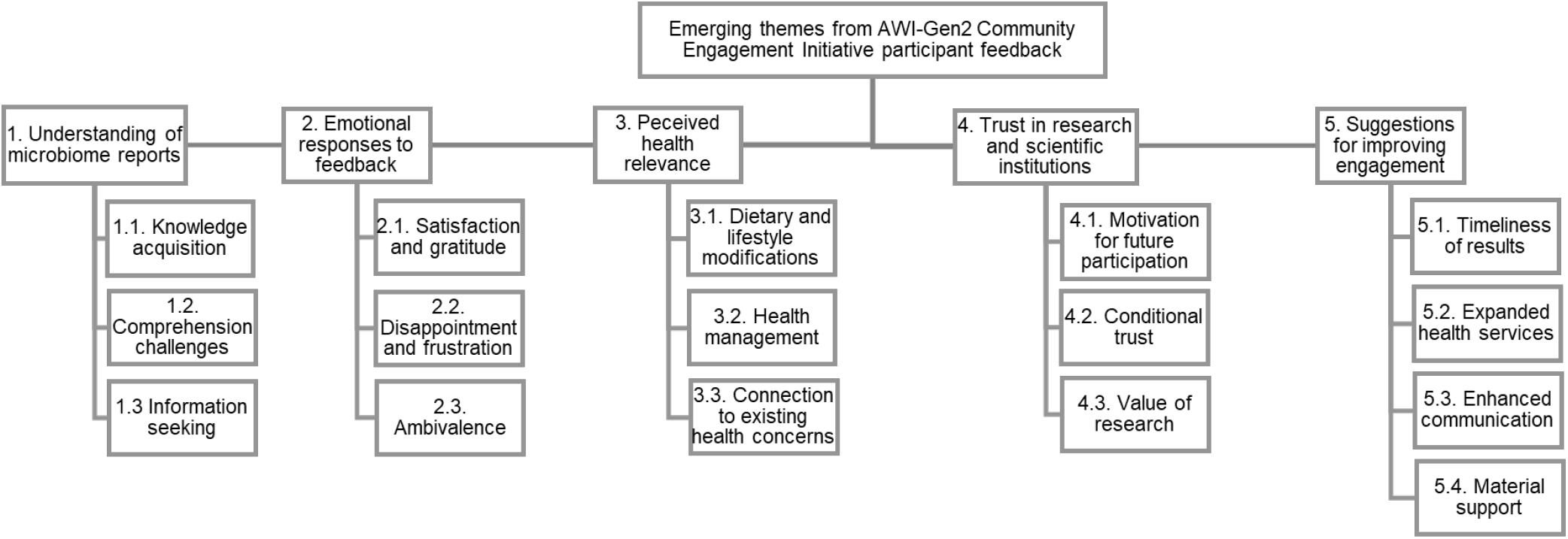
Themes and sub-themes identified through deductive and inductive analysis

### Reporting

Themes and subthemes are presented below, quantified through MAXQDA coded segments (Table S2.1) and illustrated by representative quotations provided in Supplemental File S2.

### Theme 1: Understanding of microbiome reports

The study revealed that participants’ comprehension of their microbiome results varied considerably, influencing their ability to apply the information to their lives. This theme captures how participants processed the scientific information provided, their knowledge acquisition, comprehension challenges, and their motivation to seek further information about their microbiome health.

#### Sub-theme 1.1: Knowledge acquisition

More than half of the participants (52.2%) expressed satisfaction at gaining new knowledge about their microbiome and how it affects their health. This newly found awareness represented a significant benefit of receiving their results, as participants felt empowered by understanding more about their bodies. The knowledge acquired was often described as personally meaningful and actionable.

#### Sub-theme 1.2: Comprehension challenges

Despite the value of the information provided, some participants struggled to understand the complex scientific concepts related to the microbiome. These challenges included misconceptions about bacteria being universally harmful, difficulty interpreting the meaning of their personal results, and confusion about the implications for their health. These comprehension challenges highlight the importance of accessible explanations tailored to participants’ educational backgrounds. The issue of language barriers was largely observed in the Soweto engagement, which was conducted by the researchers in English, versus the local language used in other sites.

#### Sub-theme 1.3: Information seeking

At least 20% of participants demonstrated active engagement with their results by asking specific questions about bacterial counts, health implications, and potential follow-up testing. This information-seeking behaviour reflected genuine interest in understanding their microbiome health more deeply and determining what actions they could take based on their results. They also expressed an interest in research into other diseases and health issues. Beyond comprehension, participants’ emotional responses provided insight into how feedback was received and internalized.

### Theme 2: Emotional responses to feedback

Receiving microbiome results evoked a range of emotional responses from participants, highlighting the personal significance of health information. These responses influenced participants’ overall perception of the research experience and potentially their willingness to implement health recommendations.

#### Sub-theme 2.1: Satisfaction and gratitude

The majority of participants expressed positive emotional responses upon receiving their results, including happiness, satisfaction, and gratitude. These positive emotions were directed both toward the information itself and the opportunity to learn about their health status. For many, simply receiving results after participating in research was cause for appreciation.

#### Sub-theme 2.2: Disappointment and frustration

Some participants expressed negative emotions, particularly disappointment with delayed results or unmet expectations about the scope of feedback. These feelings often stemmed from misconceptions about what the study would provide, such as expecting direct medical care rather than educational information, or frustration with the long wait time between sample collection and results delivery.

#### Sub-theme 2.3: Ambivalence

A subset of participants expressed neither strongly positive nor negative emotions about their results, instead conveying uncertainty, indifference, or mixed feelings. This ambivalence sometimes reflected confusion about the meaning or relevance of the results, or uncertainty about how to respond to the information provided. Some responses were also a reflection of whether they felt motivated to join future studies based on the engagement.

### Theme 3: Perceived health relevance

Participants varied in how they connected their microbiome results to their broader health concerns and behaviours. This theme captures the practical applications participants envisioned for their results, including dietary changes, overall health management strategies, and connections to existing health concerns.

#### Sub-theme 3.1: Dietary modifications

A prominent response to receiving microbiome results was participants’ intention to modify their diets based on the information provided. Many expressed newfound awareness of how food choices affect their gut bacteria and overall health, with specific plans to increase consumption of traditional foods, vegetables, and other health-promoting options while reducing processed foods.

#### Sub-theme 3.2: Health management

Beyond specific dietary changes, 40% of participants recognized broader implications for managing their health. They described feeling empowered to take control of their well-being through lifestyle changes rather than medication alone and appreciated understanding the connection between their microbiome and overall health status.

#### Sub-theme 3.3: Connection to existing health concerns

Some participants attempted to connect their microbiome results to specific health conditions they were experiencing or had concerns about. These connections reflected participants’ efforts to integrate the new information into their existing understanding of their health and to determine if their results could explain persistent health issues.

### Theme 4: Trust in research and scientific institutions

Participants’ willingness to trust research processes and scientific institutions emerged as a significant theme, with important implications for future community engagement. This trust was reflected in attitudes toward continued participation and recognition of research benefits, though often with important conditions elaborated on below (Sub-theme 4.2).

#### Sub-theme 4.1: Motivation for future participation

Many participants (70%) expressed enthusiasm for continued involvement in research studies, citing benefits they had experienced and anticipating future advantages. This willingness to participate reflected a positive evaluation of the current research experience and trust in the research team’s intentions and methods.

#### Sub-theme 4.2: Conditional trust

While many participants expressed willingness to participate in future research, this was often contingent on specific conditions being met. These conditions commonly related to concerns about invasive procedures, particularly blood collection, and expectations for transparency and result sharing. This conditional trust was especially evident in the Agincourt community.

### Sub-theme 4.3: Value of research

Some participants recognized the broader value of research beyond immediate personal benefits, acknowledging contributions to community health knowledge and scientific progress. This appreciation for research’s wider significance reflected a more sophisticated understanding of the research enterprise and its potential long-term benefits.

### Theme 5: Suggestions for improving engagement

Participants provided numerous recommendations for enhancing research engagement, reflecting their desire for research that better meets community needs and expectations. These suggestions covered result delivery timing, service scope, communication strategies, and material support.

#### Sub-theme 5.1: Timeliness of results

A consistent recommendation across all sites was the importance of delivering results promptly. Participants emphasised that long delays between sample collection and results feedback diminished the value of participation and eroded trust. Timely result sharing was seen as a basic requirement for respectful research engagement.

#### Sub-theme 5.2: Expanded health services

Many participants suggested that research should include broader health services beyond just microbiome testing. These recommendations included screening for conditions like cancer, providing support for women’s health issues, and offering more comprehensive healthcare access. These suggestions reflected community health priorities and expectations for reciprocity in research relationships.

#### Sub-theme 5.3: Enhanced communication

Participants valued clear, accessible communication about research processes and results. They suggested improvements to how information is shared, emphasizing the importance of language that participants can understand and materials they can reference later. Many (50%) also noted the value of being able to share knowledge with family and community members.

#### Sub-theme 5.4: Material support

Some participants expressed expectations for tangible benefits or compensation for their research participation. While views on appropriate compensation varied, many appreciated receiving vouchers, small payments, or other material support in recognition of their time and contribution. This was particularly evident in Agincourt.

### Budget and resource allocation

The allocation of resources and budget varied considerably across the three study sites, Nairobi, Agincourt, and Soweto, reflecting the unique logistical, cultural, and infrastructural needs of each setting (Table 1).

**TABLE 1.** Budget allocation for microbiome results dissemination by study site.

| Budget category | Nairobi | Agincourt | Soweto |
| --- | --- | --- | --- |
|  | (USD) | (USD)<sup>5</sup> | (USD)<sup>6</sup> |
| Personnel<sup>1</sup> | 5,147 | 3,357 | 2,507 |
| Transport<sup>2</sup> | 886 | 4,018 | 1,538 |
| Venue & overheads<sup>3</sup> | 2,302 | 6,230 | 1,015 |
| Communication & materials<sup>4</sup> | 34 | 672 | 56 |
| Total | 8,369 | 14,277 | 5,116 |

<sup>1</sup> Personnel includes research assistants, fieldworkers, administrative staff, etc.
<sup>2</sup> Transport covers participant travel reimbursements, vehicle hire, and fuel.
<sup>3</sup> Venue & Overheads includes hall/room hire, refreshments, routine site overheads, and utilities.
<sup>4</sup> Communication & Materials captures airtime, printing, transcription, and small equipment.
<sup>5</sup> May 2024, USD to ZAR = 15.93
<sup>6</sup> July 2024, USD to ZAR = 17.95

Dissemination costs varied by site, reflecting distinct logistical strategies and engagement models. Agincourt incurred the highest total budget (USD 14,277) but the lowest cost per participant (≈ USD 29). This budget was driven primarily by venue and overhead costs (USD 6,230) for community gatherings and high transportation expenses (USD 4,018), necessitated by the dispersed rural population. Personnel costs (USD 3,357) funded local field workers to ensure cultural relevance, while communication materials (USD 672) were prioritized for clarity.

In contrast, Soweto had the lowest total budget (USD 5,116) yet the highest cost per participant (≈ USD 59). Personnel (USD 2,507) was the dominant expense, focusing on research assistance for coordination, followed by transportation (USD 1,538) and venue overheads (USD 1,015). Spending on communication materials was negligible (USD 56), indicating a reliance on verbal feedback.

Nairobi occupied the middle ground with a total budget of USD 8,369 (≈ USD 43 per participant). Unlike the other sites, Nairobi’s costs were heavily skewed toward personnel (USD 5,147) to support an intensive, personalized feedback approach. Venue costs (USD 2,302) facilitated small group sessions, while transportation (USD 886) was significantly lower than Agincourt, reflecting the comparative ease of urban mobility.

## DISCUSSION

This study evaluated how participants from diverse urban and rural communities in Agincourt, Soweto (South Africa), and Nairobi (Kenya) responded to personalized gut microbiome feedback and what value and understanding they attributed to it. Our findings emphasize the considerable value participants placed on personalized and culturally relevant feedback, enhancing their understanding of microbiome health and motivating health-promoting behavioural changes. These findings demonstrate that even in low-resource contexts, metagenomic results can be translated into relatable concepts such as gut balance and dietary diversity. These outcomes align with previous evidence highlighting the positive impacts of individualized research result dissemination on health awareness and behavioural change [20].

Unlike online-first return models such as the American Gut Project and Isala, many participants in our settings had limited literacy and digital access. Feedback was therefore delivered offline through in-person sessions, visual aids, and simple language, making the educational component integral to both engagement and scientific communication rather than an optional add-on [21, 22, 5]. Our findings substantiate prior arguments that participatory microbiome initiatives must balance democratization of science with responsible interpretation and communication of uncertain data [6, 2]. Ensuring accurate comprehension is critical, as misinterpretation of compositional microbiome data could distort public perceptions of what constitutes a “healthy” microbiome.

Participants consistently highlighted the benefits of culturally grounded explanations, local-language translation, and contextualized metaphors, which substantially improved comprehension. However, comprehension was comparatively limited in Soweto, likely because verbal discussions were predominantly conducted in English – a choice necessitated by the region’s linguistic diversity and the researchers’ own language proficiency. Although the microbiome results were translated into local languages, this translation was limited to written formats, potentially limiting verbal comprehension and interactive discussion that could enhance deeper understanding. This aligns with prior findings indicating that researchers may inadvertently prioritize academic dissemination over tailored communication approaches, neglecting the importance of fully localized and interactive dissemination strategies [23]. Given the limited internet access, verbal explanation in local languages, facilitator-led Q&A, and take-home printouts were not just preferred but necessary; this contrasts with email portal returns used in American Gut and Isala and underscores the need for offline-first delivery in African contexts. Ensuring accurate comprehension is critical, as misinterpretation of compositional microbiome data could distort public perceptions of what constitutes a “healthy” microbiome. Thus, future engagements should prioritize verbal explanations and dialogue in local languages alongside written translations to ensure equitable understanding and participation

Emotional responses to the results – ranging from gratitude to frustration – revealed both the affective dimension of receiving individualized feedback and the expectations participants held about research responsiveness. Negative reactions, particularly frustration over delays, align with reports that untimely result return diminishes trust and perceived research value [13, 23]. Timely dissemination is therefore essential to maintaining trust and engagement. It is clear that there are high expectations of feedback (we had not promised personalised feedback when participants were recruited). While COVID-related pauses in the study and logistical and analysis constraints – especially the need for in-person return due to limited connectivity – posed challenges, grouping feedback into scheduled community sessions mitigated delay-related concerns. This approach differed from the online rolling returns in American Gut and Isala studies [5, 21].

Participants also described tangible health benefits from personalized feedback, translating into actionable lifestyle and dietary changes aimed at enhancing gut microbiome diversity. The motivational effect of personalized results is consistent with prior research showing that individual feedback fosters relevance and empowerment [24]. Participants from Soweto that had telephonic interviews following FDGs reported increased vegetable intake, fermented foods, and reduced processed foods changes. To make guidance tangible where budgets are constrained, we emphasized diverse, locally available plant foods and fermented staples; this parallels the “more plant types per week” message observed in American Gut but was translated into affordable, market-specific examples [5]. Given the limited emphasis on preventive care within under-resourced health systems [25, 26, 27], microbiome feedback offers a feasible preventive strategy by linking scientific findings to affordable, locally available dietary options.

Trust was a pivotal determinant of willingness to participate in future research. Although some frustrations reflected general expectations of biomedical studies rather than the microbiome sub-study, the central role of reciprocity and transparency in sustaining trust was evident. This aligns with relational ethics frameworks that emphasize accountability and cultural grounding in microbiome research [9, 10]. Concerns about invasive procedures such as blood collection highlighted the need for clear explanations of scientific purpose and specimen handling to maintain comfort and willingness [13]. In our setting, demonstrations with sample kits, simple flow diagrams, and repeated verbal consent checks effectively reinforced understanding and reduced apprehension, complementing paper consent forms.

Participant perceptions also underscored the value of feedback itself as an act of reciprocity. Participants had previously received general AWI-Gen feedback but not individualized microbiome results. Positive reactions to receiving personalized reports – and complaints about delays – illustrate the importance of closing the loop between sample collection and result sharing. Achieving this is far more complex than with routine clinical diagnostics, as microbiome results rely on multi-step workflows involving sample processing, sequencing, and intricate bioinformatic analysis rather than immediate readouts (a timeline further extended in this study by COVID-19 disruptions). Since most participants lacked internet access, grouping feedback into predictable local meetings with printed summaries managed expectations and maintained trust, representing a contextual adaptation beyond online dashboards used elsewhere. This demonstrates that even non-clinical microbiome feedback can recalibrate community expectations of research accountability and strengthen the researcher–participant relationship.

Participant recommendations for future research emphasized integrating broader health screenings with feedback delivery, particularly for cancer, hypertension, and diabetes. Such community-driven expectations echo longstanding engagement principles in African genomics that highlight reciprocity and capacity-building as foundations of ethical research [4, 28, 29]. Integrating gut-health feedback with broader screenings could position microbiome data as an entry point for holistic chronic disease prevention and promote perceptions of fairness and shared benefit [30, 31]. In low-connectivity areas, co-locating simple screenings with report-back sessions was viewed as efficient and respectful of time and travel costs.

Logistical and budgetary analyses highlighted the importance of context-specific resource allocation. Urban challenges in Soweto – such as tracking participants – and rural transportation demands in Agincourt both underscore the need for strategic planning at the inception of research. Equitable resource distribution also influences data validity, since disparities in engagement can affect sample completeness and retention across microbiome cohorts. Budgeting for print materials, language translation, and SMS or voice-call communication – rather than web-based tools – proved essential for inclusive dissemination in low-connectivity settings.

In closing, individualized gut microbiome reports, though not clinically diagnostic, can deliver meaningful value when returned transparently, accessibly, and within cultural context. By presenting results as educational tools supporting dietary diversity and fermented-food choices, we aligned participant expectations with the state of microbiome science and avoided over-interpretation. This position aligns with clinical guidance cautioning against direct use of at-home microbiome tests for health management due to the absence of reference standards [32]. It also echoes reviews noting that while diet influences microbial composition, heterogeneity in study designs limits translation and calls for harmonized analytical approaches [33]. As multi-omics analyses expand across African cohorts, embedding these community-centered feedback frameworks will ensure responsible and inclusive translation of emerging microbiome and metabolomic data. Against this backdrop, the ethical gains we document – transparent processes, culturally anchored communication, and timely reciprocity – are as important as the scientific ones, strengthening trust and sustained participation while ensuring that early returns of complex -omics data inform, empower, and do no harm.

### Limitations

This qualitative study, though robust in its triangulation and reflexivity, is not without limitations. Within the COREQ framework, we acknowledge that qualitative interpretation is inherently subjective despite safeguards for rigor and transparency. Our themes are contextually grounded; this specificity strengthens acceptability but limits generalizability. Interpretations of gut health also reflected local explanatory models and varying familiarity with biomedical microbiome concepts, which may influence cross-site comparison. Only three sites were included, some invited participants did not attend, and transcripts were not returned for member checking due to resource constraints. The feedback model was tailored to language, literacy, connectivity, and health-system realities and is therefore not a universal template. Replication elsewhere will require co-design of metaphors, visuals, facilitation language, and logistics to ensure contextual relevance. Ultimately, microbiome feedback is not solely a translational or educational act but a relational one – an exchange that constructs trust and shared meaning between scientists and communities [1, 9].

## CONCLUSION

Ultimately, this study demonstrates that returning microbiome data is not merely a logistical task but an ethical imperative that can catalyse health literacy even in the absence of clinical diagnoses. Moving beyond ad-hoc methods, the field requires a standardized, scalable framework that anchors dissemination in local realities, prioritizing verbal and visual communication in local languages over text-heavy reports. Future research must focus on longitudinal assessments to determine if such ethical engagement models sustain long-term behavioural changes and how they can be adapted for complex multi-omics data. By shifting the focus from data extraction to a structured, culturally responsive partnership, researchers can ensure that the expansion of microbiome science across Africa not only advances global knowledge but concretely benefits the communities providing the data.

## ACKNOWLEDGMENTS

We thank the AWI-Gen 2 study participants for their invaluable participation and feedback. We greatly acknowledge the APHRC, MRC/ Wits DPHRU and MRC/Wits Rural HDSS for their assistance with the study. Research reported in this publication was supported by the Fogarty International Center and National Institute of Biomedical Imaging and Bioengineering (NIBIB) and OD/Office of Strategic Coordination (OSC) and Office of Data Science Strategy (ODSS) of the National Institutes of Health, under Award Number U54 TW 012077 and a Strategic Health Innovation Partnerships grant from the South African Medical Research Council. This work was also supported in part, by the Stanford Gerald J. Lieberman Fellowship and the NIH Fogarty Global Health Equity Scholars Program (NIH FIC D43TW010540) and the Sydney Brenner Charitable Trust (SBCT). The content is solely the responsibility of the authors and does not necessarily represent the official views of the funders.

## DATA AVAILABILITY STATEMENT

### ETHICS APPROVAL

Human subject research approval was obtained from University of the Witwatersrand Human Research Ethics Committee (Protocol No: M170880, M2210108) and ethics approvals were also obtained at each study centre.

## REFERENCES

[1] O’Doherty KC, Shabani M, Dove ES, Senecal K, Knoppers BM. 2016. The Human Microbiome and Public Health: Social and Ethical Considerations. The American Journal of Bioethics 16 (11):36–48. doi:10.1080/15265161.2016.1214305.

[2] Greenhough B, Lorimer J, Brown N, Dwyer C. 2020. Setting the agenda for social science research on the human microbiome. Social Science & Medicine 252:112927. doi: 10.1016/j.socscimed.2020.112927.

[3] Tluway F, Agongo G, Baloyi V, Boua P, Kisiangani I, Lingani M, Mashaba R, Mohamed S, Nonterah E, Ntimana C, Rouamba T, Mathema T, Madala S, Maghini D, Choudhury A, Crowther N, Hazelhurst S, Sengupta D, Ansah P, Choma S, Debpuur C, Gómez-Olivé F, Kahn K, Micklesfield L, Norris S, Oduro A, Sorgho H, Tindana P, Tinto H, Tollman S, Wade A, Ramsay M, as members of AWI-Gen and the H3Africa Consortium. 02 2025. Cohort Profile: Africa Wits-INDEPTH partnership for Genomic studies (AWI-Gen) in four sub-Saharan African countries. International Journal of Epidemiology 54 (1):dyae173. doi:10.1093/ije/dyae173.

[4] Tindana P, Campbell M, Marshall P, Littler K, Vincent R, Seeley J, et al. 2015. Community engagement strategies for genomic studies in Africa: a review of the literature. BMC Medical Ethics 16 (1):24. doi:10.1186/s12910-015-0010-1.

[5] McDonald D, Hyde E, Debelius JW, Morton JT, Gonzalez A, Ackermann G, Aksenov AA, Behsaz B, Brennan C, Chen Y, DeRight Goldasich L, Dorrestein PC, Dunn RR, Fahimipour AK, Gaffney J, Gilbert JA, Gogul G, Green JL, Hugenholtz P, Knight R. 2018. American Gut: an Open Platform for Citizen Science Microbiome Research. mSystems 3 (3):e00031–18. doi:10.1128/mSystems.00031-18.

[6] Debelius JW, Morton JT, Hyde ER, McDonald D, Knight R. 2016. Citizen microbiome project: Educational and citizen science perspectives for participatory microbiome research. Journal of Microbiology & Biology Education 17 (1):46. doi: 10.1128/jmbe.v17i1.1034.

[7] Lebeer S, Ahannach S, Gehrmann T, Wittouck S, Eilers T, Oerlemans E, Condori S, Dillen J, Spacova I, Vander Donck L, Masquillier C, Allonsius C, Bron P, Van Beeck W, De Backer C, Donders G, Verhoeven V. 2023. Nature Microbiology 8 (11):2183–2195. doi:10.1038/s41564-023-01500-0.

[8] Isala Research Team. 2023. Isala: A Citizen Science Project on the Vaginal Microbiome https://isala.be/en/results/. Accessed: 2025-09-10.

[9] Bader AR, Larkin B, Garrison NA, Hudson M, Bolnick DA. 2023. A relational framework for microbiome research with Indigenous communities. Nature Microbiology 8:1688–1696. doi:10.1038/s41564-023-01457-w.

[10] Mangola SM, Black J, Delaney K, et al. 2022. Ethical microbiome research with Indigenous communities. Nature Medicine 28:2255–2257. doi: 10.1038/s41591-022-02059-2.

[11] Matimba A, Ali S, Littler K, Madden E, Marshall P, McCurdy S, Nembaware V, Rodriguez L, Seeley J, Tindana P, Yakubu A, de Vries J, H3Africa Ethics and Community Engagement Working Group. 2022. Guideline for feedback of individual genetic research findings for genomics research in Africa. BMJ Global Health 7 (1):e007184. doi:10.1136/bmjgh-2021-007184.

[12] Ralefala D, Kasule M, Matshabane O, Wonkam A, Matshaba M, de Vries J. 2021. Participants’ Preferences and Reasons for Wanting Feedback of Individual Genetic Research Results From an HIV-TB Genomic Study: A Case Study From Botswana. Journal of Empirical Research on Human Research Ethics 16 (5):525–536. doi: 10.1177/15562646211043985.

[13] Mashaba RG, Ntimana CB, Makoti P, Mothapo K, Tlouyamma J, Seakamela KP. 2024. The impact of research results feedback on the lived experiences of elderly participants in the DIMAMO health demographic site: a case of AWI-Gen Participants. medRxiv doi:10.1101/2024.11.12.24316981.

[14] Yatsunenko T, Rey FE, Manary MJ, Trehan I, Dominguez-Bello MG, Contreras M, Magris M, Hidalgo G, Baldassano RN, Anokhin AP, Heath AC, Warner B, Reeder J, Kuczynski J, Caporaso JG, Lozupone CA, Lauber C, Clemente JC, Knights D, Knight R, Gordon JI. 2012. Human gut microbiome viewed across age and geography. Nature 486 (7402):222–227. doi:10.1038/nature11053.

[15] Cybulski J, Clements J, Prakash M. 2014. Foldscope: origami-based paper microscope. PloS One 9:e98781. doi:10.1371/journal.pone.0098.

[16] Maghini DG, Oduaran OH, Olubayo LAI, et al. 2025. Expanding the human gut microbiome atlas of Africa. Nature 638:718–728. doi:10.1038/s41586-024-08485-8.

[17] Wamukoya M, Kadengye D, Iddi S, Chikozho C, System D. 04 2020. The Nairobi Urban Health and Demographic Surveillance of slum dwellers, 2002 – 2019: Value, Processes, and Challenges. Global Epidemiology 2:100024. doi: 10.1016/j.gloepi.2020.100024.

[18] Tong A, Sainsbury P, Craig J. Dec 2007. Consolidated criteria for reporting qualitative research (COREQ): a 32-item checklist for interviews and focus groups. International Journal for Quality in Health Care 19 (6):349–357. doi: 10.1093/intqhc/mzm042.

[19] VERBI Software. 2021. MAXQDA 2022 [computer software]. VERBI Software, Berlin, Germany.

[20] O’Mara-Eves A, Brunton G, Oliver S, Kavanagh J, Jamal F, Thomas J. Feb 2015. The effectiveness of community engagement in public health interventions for disadvantaged groups: a meta-analysis. BMC Public Health 15 (1):129.

[21] Isala. 2025. Isala – Let’s Swab Accessed 2025-10-17.

[22] Lebeer S, Ahannach S. 2024. The Isala Sisterhood doi: 10.6084/m9.figshare.27180102.v3.

[23] Bodison S, Sankaré I, Anaya H, Booker-Vaughns J, Miller A, Williams P, et al. 2015. Engaging the Community in the Dissemination, Implementation, and Improvement of Health-Related Research. Clinical Translational Science 8 (6):814–819.

[24] DiClemente C, Marinilli A, Singh M, Bellino L. May 2001. The Role of Feedback in the Process of Health Behavior Change. American Journal of Health Behavior 25 (3):217–227.

[25] Jackson K, Kaner E, Hanratty B, Gilvarry E, Yardley L, O’Donnell A. 2024. Understanding people’s experiences of the formal health and social care system for co-occurring heavy alcohol use and depression through the lens of relational autonomy: A qualitative study. Addiction 119 (2):268–280.

[26] Stockton M, Mazinyo E, Mlanjeni L, Sweetland A, Scharf J, Nogemane K, et al. Oct 2024. Validation of screening instruments for common mental disorders and suicide risk in south African primary care settings. Journal of Affective Disorders 362:161–168.

[27] Radebe M, Moropeng M, Patrick S. 2024. Perception of healthcare workers and patients about the impact of health facility infrastructure on healthcare services in eThekwini Municipality, KwaZulu-Natal, South Africa. International Journal of Healthcare Management 0 (0):1–10.

[28] Girardi E, Sabin C, Monforte A. Sep 2007. Late diagnosis of HIV infection: epidemiological features, consequences and strategies to encourage earlier testing. Journal of Acquired Immune Deficiency Syndromes 46 (Suppl 1):S3–S8.

[29] Fleming K, Horton S, Wilson M, Atun R, DeStigter K, Flanigan J, et al. Nov 2021. The Lancet Commission on diagnostics: transforming access to diagnostics. The Lancet 398 (10315):1997–2050.

[30] Bedeker A, Nichols M, Allie T, Tamuhla T, van Heusden P, Olorunsogbon O, et al. Feb 2022. A framework for the promotion of ethical benefit sharing in health research. BMJ Global Health 7 (2):e008096.

[31] Mwaka E, Bagenda G, Sebatta D, Nabukenya S, Munabi I. Nov 2022. Benefit sharing in genomic and biobanking research in Uganda: Perceptions of researchers and research ethics committee members. Frontiers in Genetics 13. https://www.frontiersin.org/journals/genetics/articles/10.3389/fgene.2022.1037401/full.

[32] McCallum K. 2024. Should You Do a Gut Microbiome Test? Houston Methodist Blog, Accessed 2025-10-17.

[33] Lotankar M, Houttu N, Mokkala K, Laitinen K. 2024. Diet–Gut Microbiota Relations: Critical Appraisal of Evidence From Studies Using Metagenomics. Nutrition Reviews 83 (7):e1917–e1938. doi:10.1093/nutrit/nuae192.

