## Supplementary File S2 for "Participant engagement and feedback in microbiome projects: a case of AWI-Gen 2"

**Supplemental File S2**

This file presents selected verbatim quotes from participant interviews and focus groups that informed the thematic framework described in the main text in Section A and Table S2.1. In Section B, we summarize responses under the sub-theme 4.2 “Conditional trust”, showing the distribution of positive, negative/false, and neutral perceptions across sites.

**A: Illustrative participant quotes supporting thematic analysis**

The quotes are organized by study site and aligned with the major and minor themes derived from the coding analysis (see Figure 5, Table S2.1). They provide direct evidence of participants’ perspectives on their microbiome results, emotional responses, perceived health relevance, trust in research, and recommendations for future studies.

**Table S2.1 Frequency of MAXQDA coded segments of comments assigned to themes across the AWI-Gen2 cohort**

| Theme | Sub-theme | Count | % |
|---|---|---|---|
| 1 Understanding of microbiome reports | | | |
| | 1.1 Knowledge acquisition | 323 | 52,2 |
| | 1.2 Comprehension challenges | 174 | 28,1 |
| | 1.3 Information seeking | 122 | 19,7 |
| | Theme total | 619 | 28,8 |
| 2 Emotional responses to feedback | | | |
| | 2.1 Satisfaction and gratitude | 207 | 70,6 |
| | 2.2 Disappointment and frustration | 64 | 21,8 |
| | 2.3 Ambivalence | 22 | 7,5 |
| | Theme total | 293 | 13,6 |
| 3 Perceived health relevance | | | |
| | 3.1 Dietary modifications | 46 | 9,6 |
| | 3.2 Health management | 191 | 40,0 |
| | 3.3 Connection to existing health concerns | 240 | 50,3 |
| | Theme total | 477 | 22,2 |
| 4 Trust in research and scientific institutions | | | |
| | 4.1 Motivation for future participation | 380 | 69,3 |
| | 4.2 Conditional trust | 44 | 8,0 |
| | 4.3 Value of research | 124 | 22,6 |
| | Theme total | 548 | 25,5 |
| 5 Suggestions for improving engagement | | | |
| | 5.1 Timeliness of results | 18 | 8,4 |
| | 5.2 Expanded health services | 86 | 40,2 |
| | 5.3 Enhanced communication | 107 | 50,0 |
| | 5.4 Material support | 3 | 1,4 |
| | Theme total | 214 | 9,9 |
| | **Total** | **2151** | |

### Theme 1: Understanding of microbiome reports

#### Sub-theme 1.1: Knowledge acquisition:

- Nairobi

*"I feel good because I now know what foods I should eat to improve on the beneficial bacteria like eating traditional vegetables."*

*"First, I have become self-aware. I have known where I should improve on so that the amount of good bacteria can increase."*

*"So, since this program came and we got involved I feel that it has benefited me and I will implement whatever they have taught me."*

- Soweto

*"It gives me some sort of enlightenment about what is happening around me and in South Africa and other countries. you get to have more information about what is happening to the body"*

*"They give me tools to control my health."*

*"They give me a better understanding of what I should eat and avoid."*

- Agincourt

*"Now we know the number of bacteria in our body and that is a good benefit. It can be indirect but it is good."*

*"Yes, this is so much helpful even in the community. We will learn from our results on how to take care of ourselves, by eating well."*

*"Teaching studies like this made us to feel great as we are learning."*

**Sub-theme 1.2: Comprehension challenges**

- Nairobi

*"I thought that the bacteria we have are worms and nothing else."*

*"But the medicines we do swallow have strong bacteria. We even do swallow tablets."*

*"In my opinion bacteria are all harmful. There are no beneficial bacteria because if you have it in the body all you get are problems."*

- Soweto

*"Ah, what can I say, I didn't quite understand the session. Personally, I didn't understand, I would be lying. I didn't hear anything."*

*"I like learning, but I'm still confused about what the results mean."*

*"Uhm, no but just explain things in a language we can understand. We don't hear big English words."*

- Agincourt

*"Green vegetables add more blood faster than meat."*

*"I understand that they won't see my bacteria but I think if they can take me blood, they can see them."*

*"Maybe I would have gone to the clinic if you brought them in time, and tell exact what are the bad bacteria because now I won't know which ones good and which ones are bad. This number doesn't tell anything."*

**Sub-theme 1.3: Information seeking**

- Nairobi

*"My question is; you have given us the results and I can see that my number of good bacteria is low, so will you do any follow ups?"*

*"So, you have mentioned about the foods we should eat; so what are the beneficial and harmful bacteria that we should avoid or not avoid? How do the bacteria work in our stomachs?"*

*"I would also like to know about the research because you've talked about bacteria; so from your research as APHRC [African Population and Health Research Center], what is the role or what is APHRC going to do to help those with harmful bacteria so that they can have good health?"*

- Soweto

*"I want to know more, but it would help if I knew how to use the information."*

- Agincourt

*"When will you come back with this study?"*

*"Why old woman nowadays suffers with livers?"*

**Theme 2: Emotional responses to feedback**

**Sub-theme 2.1: Satisfaction and gratitude**

- Nairobi

*"Personally I have felt good because I am self-aware."*

*"I feel good because I have known what I can do to increase the beneficial bacteria."*

*"I am grateful for the results. It has been a long journey. I am grateful because sometimes one wants to have these tests but you may not have the money. So, we are grateful to APHRC for taking our samples and doing the tests so that we get to know our status. I am also grateful for the training because we always just eat for the sake of it. We don't eat according to what is needed. We should eat foods rich in nutrients so that our bodies can fight diseases better. So, I am grateful for the training because without the training you wouldn't learn."*

- Soweto

*"I was ok, it was a relief."*

*"It made me feel appreciated."*

*"I would just like to thank you for planning that session for us and giving us feedback."*

- Agincourt

*"I'm motivated by getting back the results. I'm happy."*

*"I'm very happy to get back my results, particularly knowing about my bacteria."*

*“Thanks for reminding us on how to eat. Other people are sick but they don’t check what they eat. I’m happy to see them like this. You must always do this to us and our children so that we all have healthy lifestyle.”*

**Sub-theme 2.2: Disappointment and frustration**

- Nairobi

*“It’s taken a very long time. Maybe even some people have since died. Most people of my age have died because people are suffering from diseases.”*

*“I thought you also tested for diseases.”*

- Soweto

*“Not really. I thought we’d get real medical care, not just food advice.”*

*"Honestly, I don’t even remember giving you the samples. That was years ago! How can these results still be useful to me?"*

*“They were okay, but I thought this study would replace going to the clinic. I didn’t expectjust advice about food.”*

- Agincourt

*“We don’t know what to do about them though you are saying we must drink more water, eat vegetables and fruits. With me I am eating like that but still I feel sick. My daily life is not good. Sometimes I feel better and sometimes I don’t feel well. If these bacteria were good, I would be feeling good.”*

*“I don’t know if I will say it right or in a correct way but we need to get the full information. You were supposed to tell us that you found this and that from our bodies. I understand that you found bacteria but which ones or how many that are good. Now you are saying you don’t know which ones are good or bad. How shall we know which ones are good or bad? That was your duty and come with full information.”*

*“We will participate though you took long time and we thought it is done. Do not take long if you want us to continue with participation. What if I was very sick? Maybe I needed to see a professional because of it, what would have happened? You see! If you take something, particularly the samples in human being. Bring back the results in time.”*

**Sub-theme 2.3: Ambivalence**

- Nairobi

*“I feel okay. It is not that bad.”*

*“They are neither bad nor good. I can’t say they are good or bad.”*

- Soweto

*“I don’t know, I guess it will depend on the study.”*

*"That will depend on the type of results, as some are useful whilst some are not explained well."*

*"I don't know if they benefit me at all. It's too late to change anything now."*

- Agincourt

*"I don't know. Maybe I will. This will be determined by the type of the study."*

*"I don't know now I will decide when you come back."*

**Theme 3: Perceived health relevance**

**Sub-theme 3.1: Dietary modifications**

- Nairobi

*"Personally, I have 58 and I will improve on what I wasn't doing well. If I wasn't eating vegetables, I will try and eat the vegetables. If it's on water I will also make sure that I drink water. If I wasn't exercising, then I will improve on that."*

*"Maybe they will improve their nutrition for example I didn't used to drink water but now I will try to drink more water daily. I will reduce my drinking of sodas. I do like sodas that water. So, you've trained me that sodas are not good and so I should drink water instead. You've taught us and we've learnt something."*

*"From this result I will check on my diet and lifestyle. Before I drink water I will have to know its condition; if it's not boiled I will boil it before drinking. I will also make sure that I wash the vegetables thoroughly before cutting them. I won't want to buy vegetables that are cut by the roadside so that I cook before washing them again."*

- Soweto: The responses from Soweto were retrospective and allowed observations of lifestyle changes that were made as a result of the community engagement and feedback, such as:

*"The results made me think a lot. Since then, I've cut down on oily food, and I feel lighter. My family is even joining me, especially my granddaughter."*

*"I was a little sceptical and I thought the diet changes would not really help much, but now I feel the difference."*

*"Do you remember the probiotics you mentioned outside? I got them, and wow, what a difference! My whole family is now using them – even my husband!"*

- Agincourt

*"Now I have knowledge about food diet, from now I want to change my living style."*

*"I will visit drinking cold drinks and not regularly as I do. I will learn to drink more water."*

*"I will participate in the future and I will change my eating habit."*

**Sub-theme 3.2: Health management**

- Nairobi

*“But if you can continue educating us then we will be better off because we will prevent many diseases since when someone listens to you and does whatever you tell them then they will stay healthy. So, since you started educating me, I am now doing very well.”*

*“It is beneficial because now that I have seen the results I will continue having balanced diet which include fruits and vegetables so that I can have more good bacteria and improve my immune system.”*

*“My heartbeats used to be very fast but now I have been trained on what to do and also on the foods I should eat in moderation, my heartbeats are no longer that fast.”*

- Soweto

*“The results given were very useful as they taught me a lot about my health and well-being.”*

*“It will help with teaching me how to take care of my health better; what to eat; what to be concerned about because we do not generally have access to that information.”*

*"It’s like I have the tools to improve my health without always needing medication.”*

- Agincourt

*“I’m suffering from the illness of sugar and I think I will benefit from the diet you taught me.”*

*“I benefit to know how many types of bacteria I have and I learn how to increase it to stay healthy.”*

*“I will try to eat more vegetables, and this will also benefit my health.”*

**Sub-theme 3.3: Connection to existing health concerns**

- Nairobi

*“Because when I was in my rural home what we used to take keen care of was milk because whenever something contaminated it, it would go back. And now we wonder what to do; if you drink packaged milk it has chemicals and this other one has other chemicals as well. So should we stop drinking it?”*

*“Let me ask a question; let’s say that I have been cut in the skin and I haven’t gone for tetanus injection, can the bacteria help be heal faster?”*

- Soweto

*“They were helpful, but I started thinking – could the results explain my joint pain? I’ve had it for years, and now I wonder if it’s connected to what I eat.”*

*“I wanted to ask – do you think my results explain my fatigue? I’ve been feeling tired all the time.”*

*“I wanted to ask – do you think my results explain my arthritis?”*

- Agincourt

*"Can't you help me or have a doctor that deals with eyes? I know Wits is doing lot of things. Maybe I can get help from you."*

**Theme 4: Trust in research and scientific institutions**

**Sub-theme 4.1: Motivation for future participation**

• Nairobi

*"So I am grateful for APHRC for this project and may you continue bringing us – even if they come now and tell us that they want to test my blood, I will readily accept."*

*"It has motivated us because there were many things we didn't know but we have learned about. So, if we have another study we would attend and learn more than what we know currently."*

*"Yes, you know, you have to be happy to participate in such. The first thing I have said is that you will know that you are not alone. You know, when you are many and you are asked a question you will be ready to answer. But if you are alone you will keep asking why you are being asked such a stupid question alone. So, I would also like to participate."*

- Soweto

*"Yes, they do. If these studies can help me understand things like this, I want to be involved."*

*"Yes! I feel like I'm part of something meaningful. Plus, it's good to know what's happening inside my body."*

*"Personally, I saw that it helps me better understand myself, so it is useful to join studies in the future."*

- Agincourt

*"Yes, I'm motivated because now I have results in my hands I won't refuse in any research."*

*"I will participate in the future because you helped us, and you are still helping."*

*"I'm motivated. This can lead us to participate in the future. You must give us our results to know what's happening with our bodies."*

**Sub-theme 4.2: Conditional trust**

- Soweto

*"If it's just advice, I'd rather go to the clinic where they actually help."*

*"Well, I am not motivated. I can come but these things are annoying. When I get sick, I can just go to doctors."*

- Agincourt

*"Maybe if you can come with a treatment or something new, like benefiting the community."*

*"If you were doing like this I would participate. But the problem is that you are taking us blood for each year and you don't give back the results. We are tired of that and I can't participate because of that."*

*"I don't think I will participate in the future because you are taking us more blood and you don't help us after. We become dizzy and feel pains, but you don't check us after. Getting results really motivates but taking blood, no. I can't participate to those studies anymore."*

**Sub-theme 4.3: Value of research**

- Nairobi

*"In urban areas we have a problem because you've spoken of how to keep – one way is to make sure that we maintain cleanliness especially when we are handling food."*

*"I think you have explained that very well and now it is up to us to maintain our bodies."*

*"So when you train me then I will also share the information with the others about nutrition and tell them what they need to eat. So, when you train us, don't think that it's all getting lost, they also help us sensitize people in the community which is important."*

- Soweto

*"Keep focusing on health education – it really helps us understand. We need more sessions of this nature."*

*"They were good, and now I'm using the advice to cook healthier meals for my family."*

*"You make such a difference."*

- Agincourt

*"Thanks for reminding us on how to eat. Other people are sick but they don't check what they eat. I'm happy to see them like this. You must always do this to us and our children so that we all have healthy lifestyle."*

*"This is a good thing [study] to do as you are teaching us to eat healthy."*

*"You must carry on doing your good work, because lots of people are dying because they don't know their health status."*

**Theme 5: Suggestions for improving engagement**

**Sub-theme 5.1: Timeliness of results**

- Nairobi

*"I would suggest that next time you shouldn't take long to get back because we may forget. So you should give us back the results sooner next time. That's my suggestion."*

*"Because I had even forgotten about the stool that was taken for testing. That's why I asked you when that was taken then I remembered that there was a day I came here."*

*“In case of another study and we are tested they shouldn’t take this long with the results because we even forgot about it.”*

- Soweto: In the Soweto community engagement, the time issue was two-fold, where participants expressed frustration with the agenda timing as well as the duration between the sample collection and return of results.

*“Please respect our time because we have commitments.”*

*“Waiting so long makes me feel like I was forgotten.”*

- Agincourt

*“You were supposed to bring my results within the period of three months. Now I forgot and didn’t expect them. If you delay you are making us to get worried a lot. In future if you want us to participate, give the results in time.”*

*“Now you are coming with results and you know that we waited for them long time ago.”*

*“We waited for our results long time ago and we were angry because you don’t give back results after you collected us blood or the stools you took from us.”*

**Sub-theme 5.2: Expanded health services**

- Nairobi

*“To add on to that, I think that since APHRC will be conducting research, they can also come to the society and train people. They will also ask you whether you accept or not.”*

*“Maybe one has a cervical cancer screening because they have money, what about those who cannot afford it? So, it is very important to research on cancer especially here in Koch.”*

*“Even hypertension is very common in the community because we don’t have the drugs in the health facilities. We also have diabetes. Sometimes you are given a prescription but you don’t have the money. So, we would also like you to bring us such programs.”*

- Agincourt

*“I would also like you to come with HIV test kits if you go to the field. Our children are scared to test in the clinic because the clinic staff don’t have confidentiality. If you test today, they will tell each other, and we don’t like that.”*

*“I think when you enter to our house please check us blood pressure because we are suffering about it and we don’t know and lots of people are dying about this disease.”*

*“You must come and teach us about cancer or check us at homes because we are old.”*

**Sub-theme 5.3: Enhanced communication**

- Nairobi

*“Even if I see someone with a problem, I will tell them because we also need to share the information with others. You share the information with me and then I also share with the others.”*

*“I would like that many people who have not joined this program should also come and be trained so that they also learn. I don’t want to be selfish so that I am the only one being taught at all times. Even the others should be trained.”*

*“My recommendation is that you should support us as elderly women by educating us. So you should support us by inviting us for seminars where we can learn.”*

- Soweto

*“What I can suggest is that results be made easy to understand and explained in layman’s terms.”*

*“Just keep sharing tips and advice like you did...Maybe write it out for us, as it makes a big difference.”*

*“But some things I didn’t understand because of language. My English is not too good.”*

- Agincourt

*“Though you said you don’t know which ones are good or not good. There we won’t know whether we are healthy or ill. You must improve there and come with results that have full information.”*

*“Also, you must show us all the healthy food we must buy as we heard that at shops, they are selling healthy cooking oils that we can use. We don’t know them and if you can come with the signs, we can remember when we go to buy.”*

*“You must come to our prayer sessions where we have gathered as women and teach about this.”*

**Sub-theme 5.4: Material support**

- Nairobi

*“And then I would like to know, can the program feed us even for a month then test us after that?”*

*“My question is, now that we are in this group is there something you can give us so that we can get these foods and increase the beneficial bacteria?”*

- Soweto

*“Yes, they were useful, and I liked the R150. It made me feel appreciated.”*

- Agincourt

*“If you were giving us vouchers, we would be able to buy fruits like apples and put them inside the fridge.”*

*“I’m happy because you are also giving us vouchers and blankets unlike in the past where we participated, you took us blood and give us nothing.”*

*“But you have to assist us with something, for example you have to build us small things like toilets.”*

**B. Distribution of perceptions under sub-theme 4.2: Conditional trust**

Within the sub-theme of *Conditional trust*, participant perceptions were categorized into four groups to capture the direction of responses: positive (expressing affirming thoughts or feedback toward the researchers, the study, the microbiome/bacteria, or intended behaviour after the session), negative/false (expressing doubts, critical views, or misconceptions that were inaccurate), neutral (general comments or observations not directly tied to the study, session, or microbiome findings) and unrelated (concerns regarding blood collection i.e. phlebotomy from the wider AWI-Gen study). These perceptions overlapped with “Sub-theme 1.2: Comprehension challenges”. Figure S2.1 illustrates the distribution of these responses across sites, highlighting variation in how conditional trust was expressed.

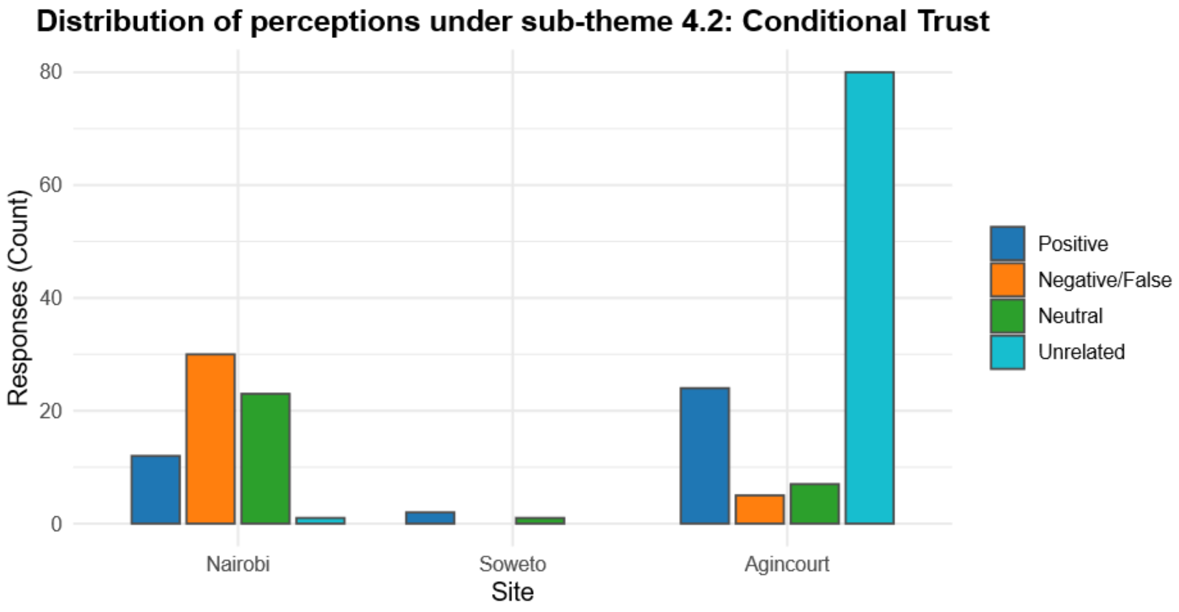


| Site | Positive | Negative/False | Neutral | Unrelated |
|---|---|---|---|---|
| Nairobi | 12 | 30 | 23 | 1 |
| Soweto | 2 | 0 | 1 | 0 |
| Agincourt | 24 | 5 | 7 | 80 |

Figure S2.1: Distribution of participant perceptions within the sub-theme 4.2 "Conditional trust," categorized as positive, negative/false, neutral, or unrelated by study site.
