## Supplementary File S1 for "Participant engagement and feedback in microbiome projects: a case of AWI-Gen 2"

**Supplemental File S1**

**Foldscope activities during microbiome feedback sessions**

This file provides a detailed description of the Foldscope activities used in community engagement sessions. A description of the preparation of Foldscope device and the prepared slides supplied in the kit is detailed below. In addition, we summarizes the application activities using water samples collected from participants' daily environment (e.g., drinking and stored water). A photographs of the Foldscope device is shown below in Figure S1

**Foldscope device:**

Foldscopes are ultra-low-cost, origami-based paper microscopes that achieve magnification up to 140× and can resolve structures as small as 2 microns. They can be assembled in minutes and used with natural light, making them highly portable and accessible in low-resource settings.

**Application in engagement sessions**

To enhance comprehension of microbiome concepts, Foldscopes were incorporated as interactive visual tools during participant feedback sessions. Participants were first introduced to prepared slides supplied with the Foldscope kit, showing common microbial structures.

During feedback sessions, participants were first introduced to prepared slides supplied with the Foldscope kit, showing common microbial structures. In addition, facilitators used samples of water collected from participants' daily environment (e.g., household drinking water and stored water) to demonstrate the presence of microbes not visible to the naked eye. This approach linked abstract microbiome concepts to participants' lived experiences, making the invisible microbial world both tangible and personally relevant.

Facilitators explained how these visible microbes differed from gut microbes, while reinforcing the broader idea of microbial diversity, its invisibility, environmental exposures, and its importance for health. Participants expressed curiosity and enthusiasm when using the Foldscopes, often commenting that it felt like "seeing the hidden world." This activity complemented the visual metaphors (gardens, soccer teams) used in group discussions and created memorable analogies that supported long-term understanding of microbiome diversity..

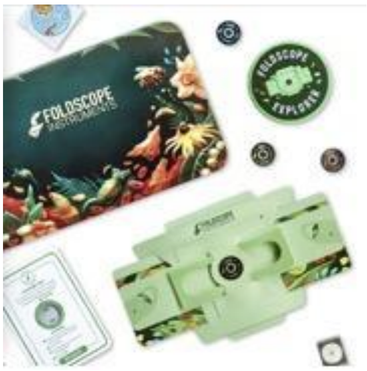


**Figure S1.** Use of Foldscopes during microbiome results feedback sessions.

Foldscope kit and components, including the paper-based microscope used in engagement sessions. The participants used the Foldscopes to observe prepared slides and water samples during a feedback session.
