## Supplementary figures and images for "Participant engagement and feedback in microbiome projects: a case of AWI-Gen 2"

### Supplementary File S1

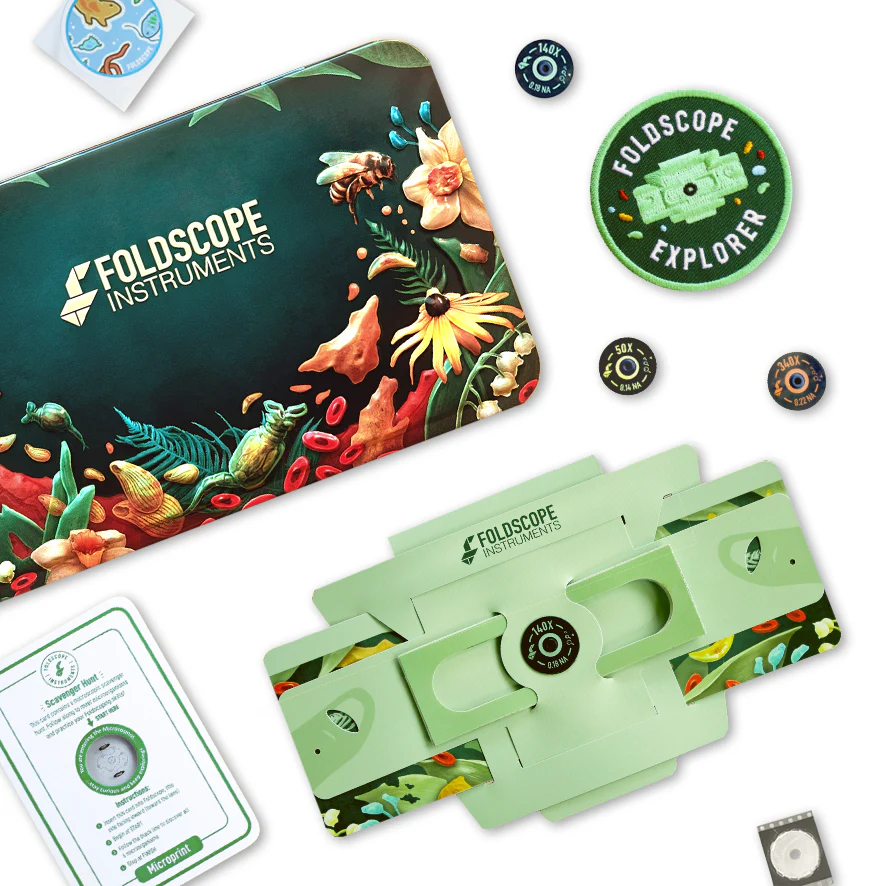
